# Contribution of odorant binding proteins to olfactory detection of (Z)-11-hexadecenal in *Helicoverpa armigera*

**DOI:** 10.1101/2020.06.10.145235

**Authors:** Hao Guo, Ping-Ping Guo, Ya-Lan Sun, Ling-Qiao Huang, Chen-Zhu Wang

**Affiliations:** State Key Laboratory of Integrated Management of Pest Insects and Rodents, Institute of Zoology, Chinese Academy of Sciences, Beijing, PR China; CAS Center for Excellence in Biotic Interactions, University of Chinese Academy of Sciences, Beijing, PR China; Forest College, Henan University of Science and Technology, Luoyang, PR China

**Author notes:** Corresponding author: Chen-Zhu Wang, State Key Laboratory of Integrated Management of Pest Insects and Rodents, Institute of Zoology, Chinese Academy of Sciences, 1 Beichen West Road, Chaoyang District, Beijing, China.

**Keywords:** Pheromone binding proteins, General odorant binding proteins, Olfactory receptors, *Helicoverpa armigera*, *Drosophila melanogaster*, Single sensillum recording

## Abstract

*Helicoverpa armigera* utilizes (Z)-11-hexadecenal (Z11-16:Ald) as its major sex pheromone component. Three pheromone binding proteins (PBPs) and two general odorant binding proteins (GOBPs) are abundantly expressed in male antennae of *H. armigera*. However, their precise roles in the olfactory detection of Z11-16:Ald remain enigmatic. To answer this question, we first synthesized the antibody against HarmOR13, a pheromone receptor (PR) primarily responding to Z11-16:Ald and mapped the local associations between PBPs / GOBPs and HarmOR13. Immunostaining showed that HarmPBPs and HarmGOBPs were localized in the supporting cells of sensilla trichodea and sensilla basiconica respectively. In particular, HarmPBP1 and HarmPBP2 were colocalized in the cells surrounding the olfactory receptor neurons (ORNs) expressing HarmOR13. Next, using two noninterfering binary expression tools, we heterologously expressed HarmPBP1, HarmPBP2 and HarmOR13 in *Drosophila* T1 sensilla to validate the functional interplay between PBPs and HarmOR13. We found that the addition of HarmPBP1 or HarmPBP2 significantly increased the sensitivity of HarmOR13 to Z11-16:Ald. However, the presence of either HarmPBP1 or HarmPBP2 was ineffective to change the tuning breadth of HarmOR13. Taken together, our results support the idea that PBPs are contributors to the peripheral olfactory sensitivity but do not affect the selectivity. Lastly, we discovered that HarmOR13 and the *Drosophila* OR67d employed a similar coding mechanism to detect pheromones, suggesting that pheromone detection across different insect orders appears to co-opt a conserved molecular principle to recognize pheromone ligands.

## Introduction

Noctuid moths rely heavily on a complex peripheral olfactory system to forage, oviposit and localize mates (Hansson, 1995). Moth antennae are studded with three major types of sensilla (Hallberg et al., 1994; Koh et al., 1995). Sensilla trichodea are specialized in pheromone detection, while sensilla basiconica and coeloconica are tuned to general odorants (Hansson and Stensmyr, 2011). The sensitivity of insect peripheral olfaction is mediated by serval class of chemosensory proteins including odorant binding proteins (OBPs), olfactory receptors (ORs) and odorant degrading enzymes (ODEs) (Guo and Wang, 2019; Leal, 2013; Su et al., 2009). Insect ORs are predicted as heterotetramers consisting of two subunits of narrowly tuned ORs and two subunits of highly conserved odorant receptor co receptor (ORco) (Butterwick et al., 2018). OBPs are a group of soluble proteins with low molecular weights of 10-14 kDa present at high concentration (up to 10 mM) in the extracellular sensillum lymph bathing the olfactory receptor neurons (ORNs) in morphologically distinct sensilla (Pelosi, 2005; Pelosi et al., 2006). The widely accepted yet debated model regarding their function is that OBPs bind odorants in the sensillar lymph and carry them to ORs on the membranes of ORNs (Gonzalez et al., 2019; Larter et al., 2016; Pelosi et al., 2018; Sun et al., 2018; Vogt et al., 1991; Xiao et al., 2019).

Pheromone binding proteins (PBPs), a subtype of OBPs, are exclusively found in sensilla trichodea and are dedicated to pheromone detection (Leal et al., 2005; Steinbrecht et al., 1995a; Vogt and Riddiford, 1981). Numerous studies have shown that the binding affinity of PBPs is tuned to species-specific pheromones (Chang et al., 2015; Guo et al., 2012; Maida et al., 2000; Pophof, 2004; Sun et al., 2013b). However, the degree of specificity of PBPs reported in different studies varies greatly. Radioactive binding assay demonstrated that ApolPBP1 of *Antheraea polyphemus* can discriminate between (E,Z)-6,11-hexadecadienyl acetate and (E,Z)-6,11-hexadecadienal (Maida et al., 2000), whereas competitive fluorescence binding assay revealed that the same protein binds these two pheromone compounds fairly well (Campanacci et al., 2001). In *Bombyx mori*, BmorPBP1 specifically interacts with bombykol to sharpen the selective response of BmorOR1 (Große-Wilde et al., 2006; Pophof, 2002), whereas only poor binding selectivity of BmorPBP1 toward bombykol and bombykal was revealed by *in vitro* fluorescence binding assay (Gräter et al., 2006). With the advent of genome editing in Lepidoptera, a handful of PBP mutants have been made in *Bombyx mori* (Shiota et al., 2018), *H. armigera* (Ye et al., 2017), *Spodoptera litura* (G.-H. Zhu et al., 2016; Zhu et al., 2019) and *Chilo suppressalis* (Dong et al., 2019). In particular, *BmorPBP1* mutant presents decreased sensitivity to both bombykol and bombykal (Shiota et al., 2018). Furthermore, the presences of PBPs could increase both sensitivity and selectivity of PRs expressed in *Xenopus* oocyte, and this effects seems to depend on the PBP-PR combination (Sun et al., 2013b, 2013a; Xu et al., 2012). Nevertheless, these contradictory findings prevent our understanding of the precise role of PBPs in pheromone detection. In addition, general odorant binding proteins (GOBPs) are also implicated in pheromone detection as increasing binding data have shown that GOBPs favorably bind to pheromones (He et al., 2010; Li et al., 2016; Liu et al., 2015; Zhang et al., 2020; Zhou et al., 2009; J. Zhu et al., 2016). In most cases, two GOBPs are found in moth antennae of both sexes generally expressed in sensilla basiconica (Li et al., 2016; Steinbrecht et al., 1995b; Wang et al., 2003), but occasionally also in sensilla trichodea (Huang et al., 2018; Jacquin-Joly et al., 2000). Apart from the binding data, there is a paucity of convincing evidence to support the role of GOBPs in pheromone detection.

*Helicoverpa armigera*, a major pest in agriculture and horticulture worldwide, utilizes (Z)-11-hexadecenal (Z11-16:Ald) as principal sex pheromone component (Wang et al., 2005). Three types of sensilla trichodea, namely type A, B and C (ratio: 84:5:11) have been functionally characterized on the antennae of males (Xu et al., 2016). Basically, type A responds specifically to Z11-16:Ald, type B to (Z)-9-tetradecenal (Z9-14:Ald), and type C to (Z)-9-hexadecenal (Z9-16:Ald), Z9-14:Ald, (Z)-11-hexadecenyl acetate (Z11-16:OAc) and (Z)-11-hexadecenol (Z11-16:OH) (Wu et al.,2013; Xu et al., 2016). *In vitro* competitive fluorescence binding assays demonstrate that both HarmPBP1 and HarmPBP2 indiscriminately bind to Z11-16:Ald, Z9-16:Ald, Z11-16:OH, (Z)-11-hexadecenol (Z9-16:OH), Z11-16:OAc and (Z)-9-hexadecenyl acetate (Z9-16:OAc), whereas HarmPBP3 binds the two acetates (Guo et al., 2012; Zhang et al., 2012). Moreover, HarmPBP1 knock-out displays diminished sensitivities to Z11-16:Ald, Z9-16:Ald and Z9-14:Ald (Ye et al., 2017), but this phenotype could be partially attributed to genetic backgrounds and off-target effects. Moreover, the roles of HarmPBP2 and HarmPBP3 have not been addressed in that study. Taken together, it is still unclear which PBPs and/or GOBPs of *H. armigera* are involved in detecting Z11-16:Ald and whether their presences affect the specificity of Z11-16:Ald detection.

*Drosophila* peripheral olfactory system has been shown to be a reliable tool to deorphanize moth ORs (Hallem et al., 2004; Fouchier et al., 2017; Ronderos et al., 2014; Ueira-Vieira et al., 2014; Wang et al., 2016) and study the functional interaction between PBPs and pheromone receptors (PRs) (Syed et al., 2006). A seminal study reports that the presence of BmorPBP considerably enhances the response of BmorOR1 to bombykol in *Drosophila* ab3 sensilla (Syed et al., 2006). Moreover, the *Drosophila* ab3A neuron is not an ideal system to express moth PRs (Syed et al., 2010). Here, we employed two noninterfering binary expression tools i.e., *GAL4*-*UAS* (Brand and Perrimon, 1993) and *LexA*-*LexAOP* (Lai and Lee, 2006), to express HarmPBPs in the *Drosophila* T1 sensillum lymph and HarmOR13 in the *Drosophila* OR67d ORN, respectively. Using this system in combination with immunofluorescence staining, we investigated the distribution patterns of HarmPBPs and HarmGOBPs and their physical connections to HarmOR13, and finally defined the role of HarmPBP1 and HarmPBP2 in the detection of Z11-16:Ald.

## Results

### Spatial distribution of HarmPBPs and HarmGOBPs on male antennae of *H. armigera*

Since both PBPs and GOBPs are implicated in binding pheromones, we expect to find both types of OBPs in the sensilla trichodea of males. To verify this hypothesis, we raised polyclonal antibody against HarmPBPs and HarmGOBPs from different hosts and performed immunofluorescent staining to map HarmPBPs and HarmGOBPs on antennae of males. Figure 1 shows that the localizations of HarmPBPs and HarmGOBPs were mutually exclusive and co-localization was never found in the longitudinal sections (Fig. 1b-f). HarmPBPs were only detected in the auxiliary cells underneath the sensilla trichodea, while the expression of HarmPBPs were stricktly restricted to the cells associated with the sensilla basiconica (Figure. 1b-f). This distribution pattern therefore suggests that HarmPBPs are specifically involved in pheromone detection while HarmGOBPs are involved in smelling general odorants.

**Figure 1.**
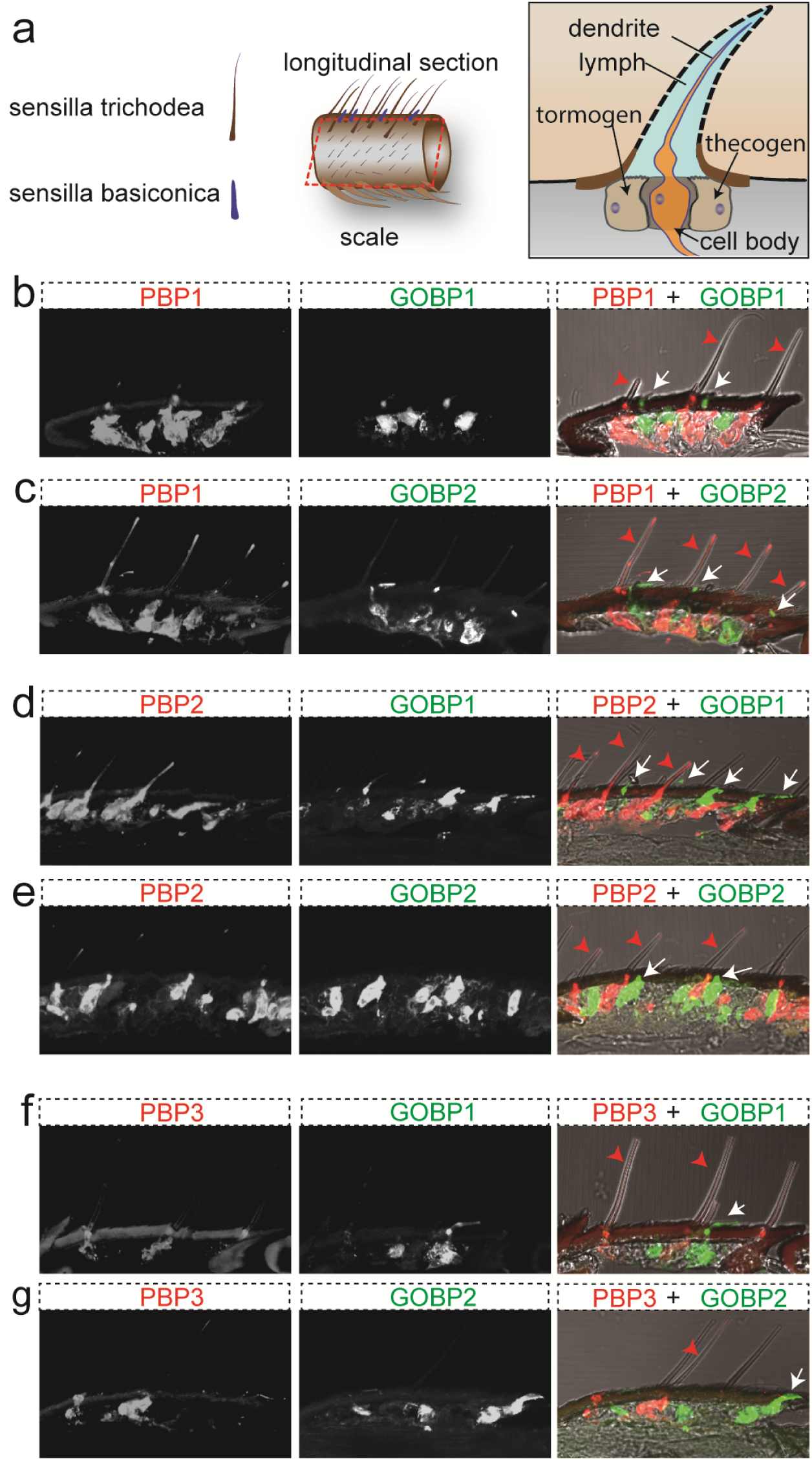
Immunolocalization of HarmPBPs and HarmGOBPs on antennae of males. HarmPBPs are typically associated with sensilla trichodea and HarmGOBPs with sensilla basiconica. The red arrow heads point to sensilla trichodea and the white arrows point to the sensilla basiconica. **(a)** left panel: the morphology of sensilla trichodea and sensilla basiconica. middle panel: scheme of the preparation of longitudinal sections. right panel: classic schematic draw of a chemosensillum trichodea. OBPs are synthesized in the tormogen cells and thecogen cells and then secreted into the sensillum lymph bathing ORNs. **(b-f)** distribution of the three HarmPBPs and the two HarmGOBPs. Scale bars: 20 µm.

Next, we asked whether members of HarmPBPs or HarmGOBPs were colocalized in the same sensillum. The immunostaining results showed that HarmPBP1and HarmPBP2 were always co-localized (Fig. 2a). Intriguingly, a small fraction of HarmPBP1 positive cells also expressed HarmPBP3 (Fig. 2b). Taken together, these results identify two categories of PBP positive cells: category A expressing HarmPBP1 & HarmPBP2 and category B harboring all three PBPs. We counted the number of category B cells on sections and found that, on average, HamPBP3 expressing cells accounted for about 18.7% of all PBP positive cells (Figure 2c), which reflects the distribution of the B and the C type of sensilla trichodea on the antennae of males. In addition, HarmGOBP1 and HarmGPBP2 were co-localized in the sensilla basiconica (Figure 2d).

**Figure 2.**
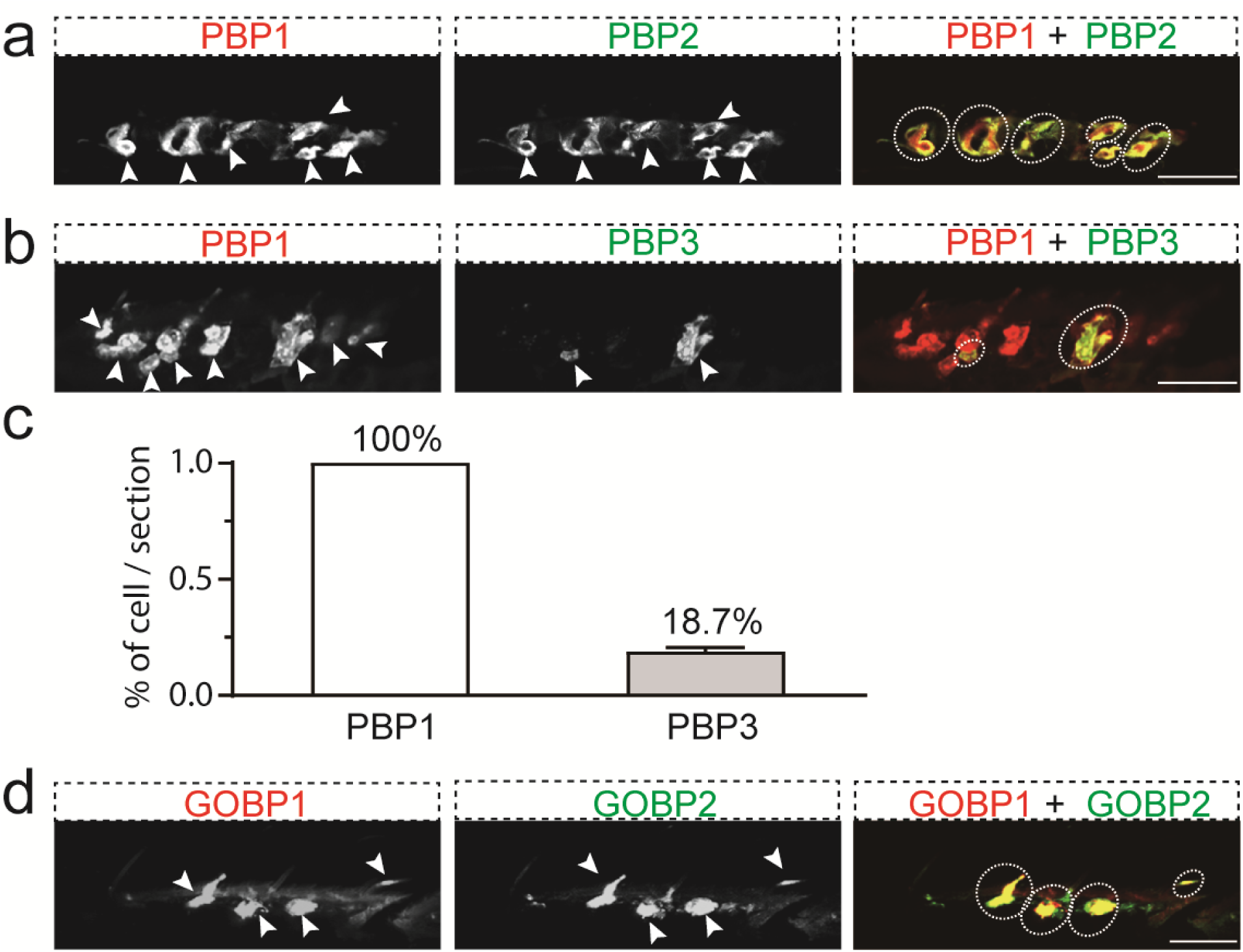
Colocalization of HarmPBPs and HarmGOBPs. **(a)** HarmPBP1 and HarmPBP2 are always expressed in the same cells. **(b)** HarmPBP1 and HarmPBP3 are erratically colocalized. **(c)** quantification of the number of cells positive for HarmPBP1 and HarmPBP3. **(d)** HarmGOBP1 and HarmGOBP2 are colocalized in the same cells. Scale bars: 20 µm.

### Spatial relationship of HarmOR13 to HarmPBPs and HarmGOBPs

To investigate the physical association of HarmOR13 to both types of OBPs, we produced a mouse polyclonal antiserum against HarmOR13, the receptor responsive to Z11-16:Ald. We observed that HarmPBP1 and HarmPBP2 expressing cells were closely located to HarmOR13 positive ORNs (Fig. 3a, b), whereas the cells additionally expressing HarmPBP3 were not (Fig. 3c). It is noteworthy that a small portion of HarmPBP1 and HarmPBP2 positive cells were not associated with HarmOR13 ORNs (Fig. 3a, b), suggesting their existence in sensilla trichodea other than the type A. The localization on horizontal sections also exhibited the same pattern (Fig. 4). Moreover, RNA *in situ* hybridization results confirmed the immunofluorescence results (Supple. e fig. 1). In addition, neither GOBP1 nor GOBP2 expressing cells showed spatial relationships with HarmOR13 ORNs (Fig. 3d, e), which was in line with their confined presence to sensilla basiconica responding to general odorants (Fig. 1d-f). In summary, the localization patterns reported here supports the involvement of HarmPBP1 and HarmPBP2 in the detection of Z11-16:Ald, and concurrently rules out the participation of HarmPBP3, HarmGOBP1 and HarmGOBP2 in this process.

**Figure 3.**
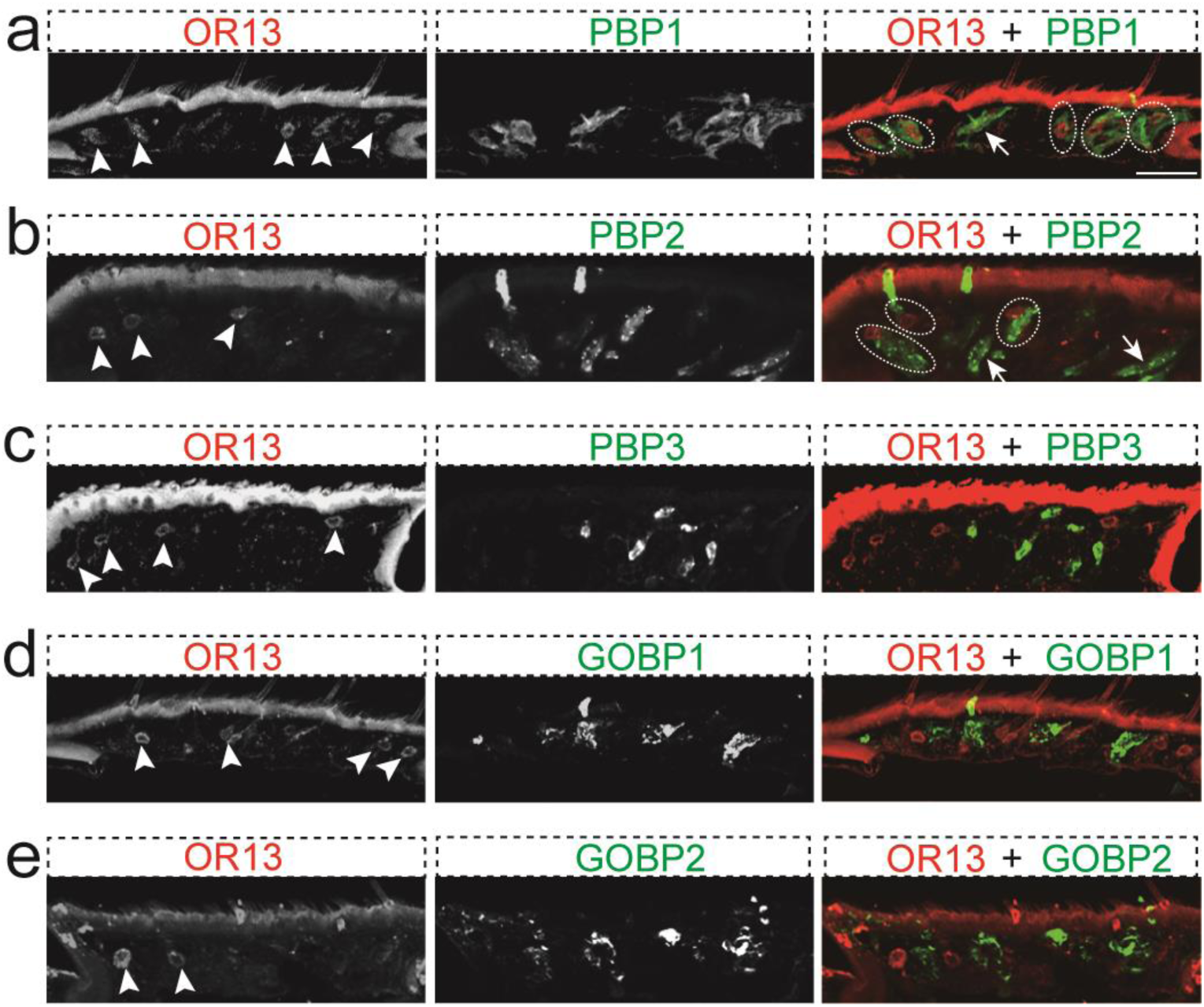
Association of HarmOR13 with HarmPBPs and HarmGOBPs. The expression of HarmOR13 is pointed by white arrow heads. The white dashed circles highlight adjacent localizations. **(a)** adjacent localization of HarmOR13 and HarmPBP1. The arrow points to the support cell that are not localized with HarmOR13 in the same sensilla. **(b)** adjacent localization of HarmOR13 and HarmPBP2. The arrows point to the support cells that are not localized with HarmOR13 in the same sensilla. **(c)** localization of HarmOR13 and HarmPBP3. **(d)** localization of HarmOR13 and HarmGOBP1. **(e)** localization of HarmOR13 and HarmGOBP2. Scale bars: 20 µm.

**Figure 4.**
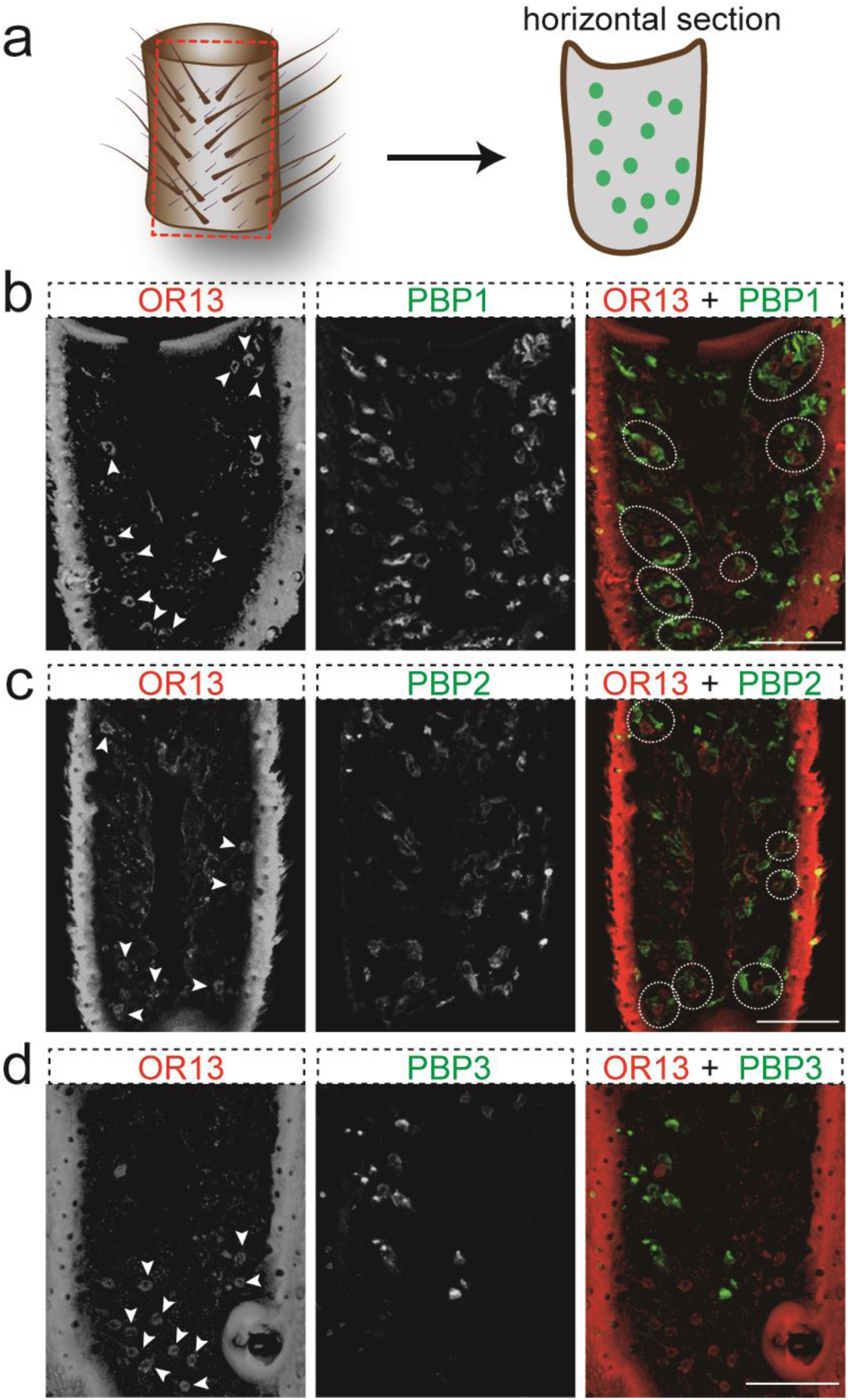
Localization of HarmOR13 and HarmPBPs on horizontal section. Close localizations are highlighted by dash circles. **(a)** localization of HarmOR13 and HarmPBP1 revealing close associations of HarmPBP1 positive supporting cells with HarmOR13 ORNs. **(b)** localization of HarmOR13 expressing ORNs and HarmPBP2 supporting cells showing their presences in the same sensilla. **(c)** localization of HarmOR13 expressing ORNs and HarmPBP3 supporting cells indicating their presences in distinct sensilla. Scale bars: 20 µm.

### Heterologous expression of HarmPBP1, HarmPBP2 and HarmOR13 to *Drosophila* T1 sensilla

Having demonstrated that HarmPBP1 and HarmPBP2 expressing cells are closely located to HarmOR13 ORNs, we went on to probing the roles of HarmPBP1 and HamPBP2 in Z11-16:Ald detection. We employed two binary expression tools, i.e., *GAL4*-*UAS* and *LexA*-*LexAOP*, to ectopically express HarmPBPs and HarmOR13 in *Drosophila* T1 sensilla (Fig. 5a). Using *LUSH-GAL4*, we successfully expressed HarmPBP1 and HarmPBP2 in the lymph of *Drosophila* T1 sensilla through supporting cells secretory pathway (Fig. 5b). In parallel, HarmOR13 was expressed in the ORNs by the driver line *ORco-LexA* and the receptor was fully functional as revealed by its response to Z11-16:Ald sensitivity in OR67d ORN (Fig. 5c).

**Figure 5.**
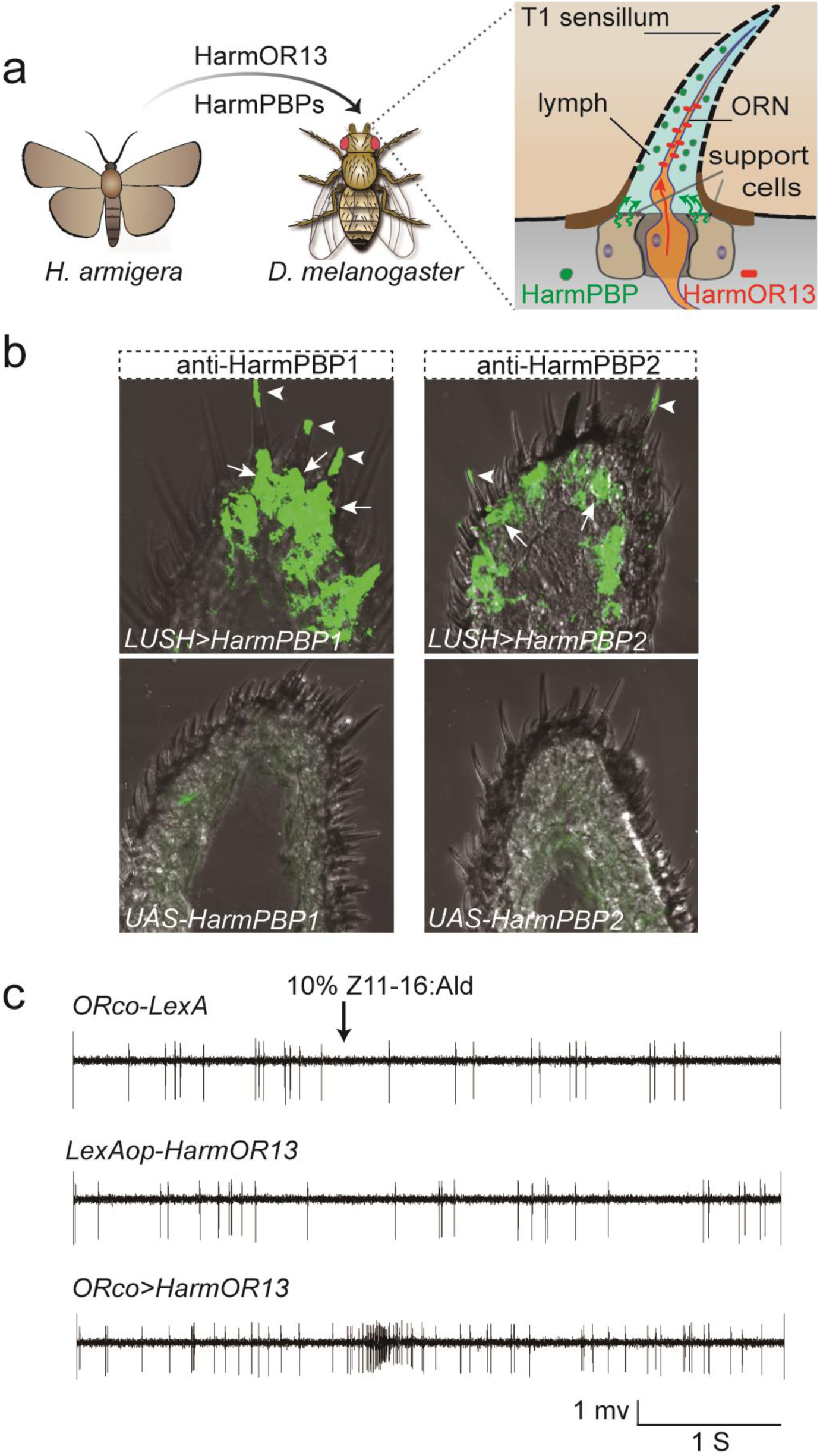
Heterologous expression of HarmPBP1, HarmPBP2 and HarmOR13 in *Drosophila* T1 sensilla. **(a)** schematic drawing illustrating the expression of HarmPBPs and HarmOR13 in *Drosophila* T1 sensillum lymph and ORN respectively. **(b)** Expression of HarmPBP1 (left panel) (genotype: *UAS-HarmPBP1/CyO;LUSH-GAL4/TM6B*) and HarmPBP2 (right panel) (genotype: *UAS-HarmPBP2/CyO;LUSH-GAL4/TM6B*) to the sensillum lymph through the supporting cell secretory pathway. No HarmPBPs were detected in the UAS control flies (lower two panels) by immunofluorescence staining (genotype: *UAS-HarmPBP1/CyO; Sb/Tm6B* and *UAS-HarmPBP2/CyO; Sb/TM6B*). The white arrows point to the supporting cells expressing HarmPBPs and the arrow heads point to the HarmPBPs secreted in the sensillum lymph. **(c)** representative traces showing responses to 10% Z11-16:Ald. (genotype: *Sp/CyO; Orco-LexA* and *LexAOP-HarmOR13;Sb/TM6B* and *LexAOP-HarmOR13/*CyO; *Orco-LexA/TM6B*)

### Functional interactions between HarmPBP1 / HarmPBP2 and HarmOR13

To determine which PBP was functionally correlated with HarmOR13, we compared the response spectra of HarmOR13 to a panel of 29 compounds including the pheromones from several moth species with and without HarmPBP1 / HarmPBP2. With a wide panel of ligands, we could define with greater accuracy the receptor specificity. First, in the absence of native HarmPBP1, HarmOR13 expressed in the *Drosophila* OR67d ORNs was narrowly tuned to the pheromone Z11-16:Ald and a structural analog, Z11-14:Ald (Fig. 6a). Next, we observed that the presence of HarmPBP1 could increase the sensitivity to Z11-16:Ald by 89% and the sensitivity to Z11-14:Ald by 77% (Fig. 6b). However, these increases in the receptor sensitivity to both compounds are not statistically significant (two-way ANOVA F_(1,16)_ = 0.34. *P* = 0.28) (Fig. 6b). This fact is clear from the response curves showing the observable increase starting from the 3% Z11-16:Ald dose (Fig. 6c). The presence of HarmPBP2 also increased the sensitivity of the ORN housing HarmOR13 to both Z11-16:Ald and Z11-14:Ald without alter its selectivity (Fig. 6d-f). Two-way ANOVA analysis indicated that the increase of sensitivity to 10% Z11-16:Ald by HarmPBP1 and HarmPBP2 was not statistically different between the two PBPs (F_(1,16)_ = 0.34. *P* = 0.57), implying HarmPBP1 and HarmPBP2 have comparable binding affinities to Z11-16:Ald and are functioning in a similar manner.

**Figure 6.**
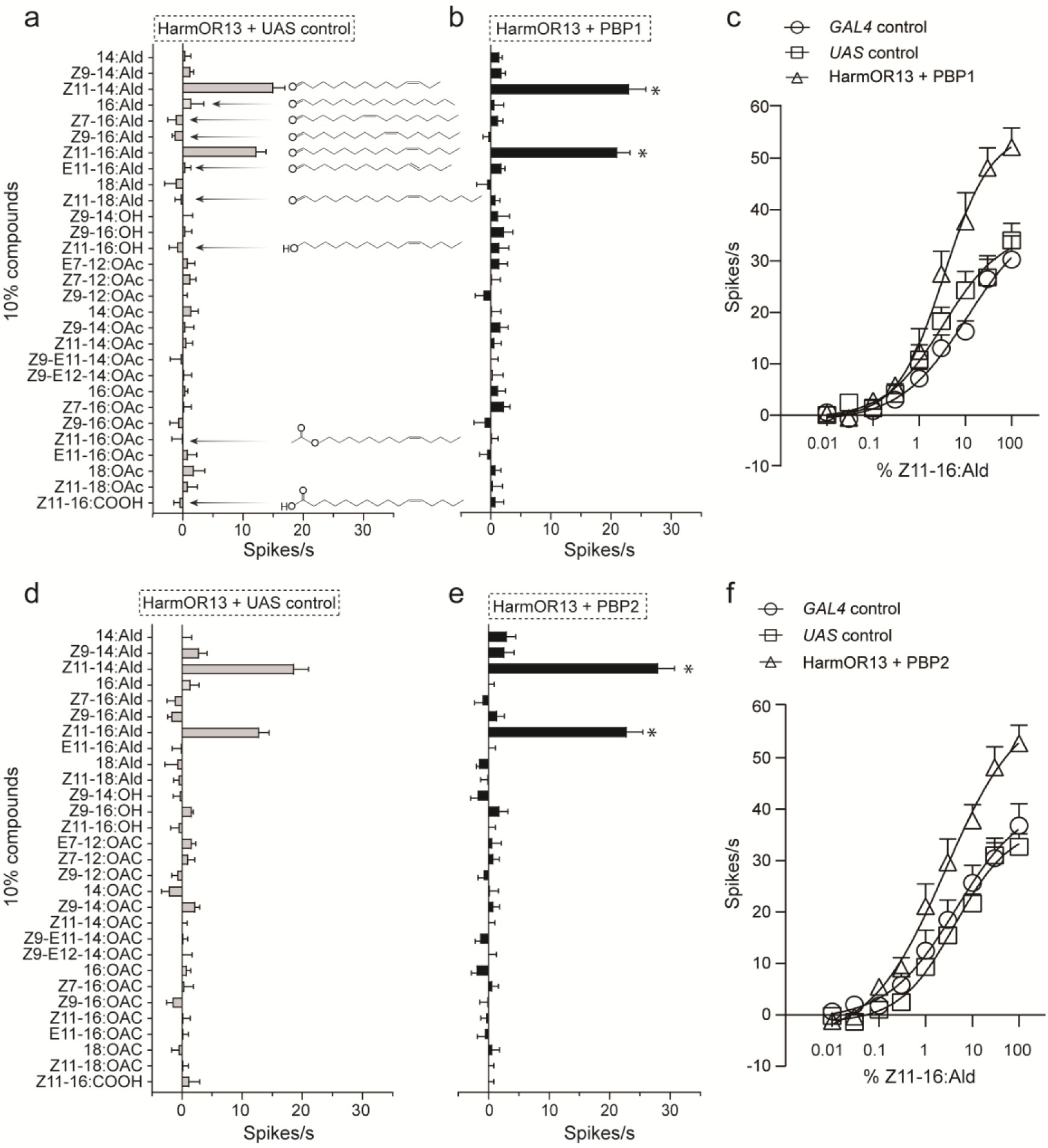
Response spectra of HarmOR13 with and without HarmPBPs. **(a)** response spectrum of HarmOR13 in the absence of native HarmPBP1 (left panel) (genotype: *LexAOP-HarmOR13/UAS-HarmPBP1; ORco-LexA/TM6B*). The T1 sensilla expressing HarmOR13 are challenged with 30 µl of 10% compounds diluted in paraffin oil. **(b)** response spectrum of HarmOR13 in the presence of HarmPBP1 (genotype: *LexAOP-HarmOR13/UAS-HarmPBP1; ORco-LexA/LUSH-GAL4*). **(c)** dose response of HarmOR13 with and without HarmPBP1 to Z11-16:Ald (GAL4 genotype: *LexAOP-HarmOR13/CyO; ORco-LexA/LUSH-GAL4*). The doses used are 30 µl of 0.01%, 0.03%, 0.1%, 0.3%, 1%, 3%, 10%, 30% and 100% Z11-16:Ald diluted in paraffin oil. **(d)** response spectrum of HarmOR13 in the absence of native HarmPBP2 (left panel) (genotype: *LexAOP-HarmOR13/UAS-HarmPBP2; ORco-LexA/TM6B*). **(e)** response spectrum of HarmOR13 in the presence of HarmPBP2 (genotype: *LexAOP-HarmOR13/UAS-HarmPBP2; ORco-LexA/LUSH-GAL4*). **(f)** The dose response of HarmOR13 in the absence and presence of HamPBP2 to Z11-16:Ald (GAL4 genotype: *LexAOP-HarmOR13/CyO; ORco-LexA/LUSH-GAL4*). Data are plotted with ± SEM. N=5 for each recording. Asterisks* indicate *P* < 0.05.

### The molecular principle of ligand-receptor coupling

We serendipitously found that HarmOR13 expressed in the *Drosophila* T1 sensilla not only responded to Z11-16:Ald but also exhibited even higher sensitivity to Z11-14:Ald which is two carbon atoms shorter (Fig. 6a). This revealed a flexibility of HarmOR13 to accommodate pheromones with different lengths. Intriguingly, HarmOR13 did not respond to a longer structural analog, Z11-18:Ald (Fig. 6a), indicating that there is a limit to the lengths of pheromone molecules. Moreover, HarmOR13 did not show any responses to 16:Ald, Z9-16:Ald and Z7-16:Ald (Fig. 6a), implying the necessity of the double bond as well as its position for recognition by this PR. Furthermore, the insensitivity of HarmOR13 to E11-16:Ald suggests that the geometric isomerism is a key factor to mediate the recognition by PRs (Figure 6a). In addition, Z11-16:OAc, Z11-16:OH and Z11-16:COOH, which share the same backbone of Z11-16:Ald but differ in the polar heads, were ineffective in activating HarmOR13 (Figure 6a), showing the importance of the functional group for the pheromone-receptor interaction. Next, we wondered whether similar criteria of chemical recognition could also be applied to the PRs of *Drosophila*. To this end, we challenged the OR67d ORN with the panel of 29 compounds. We observed that OR67d selectively responded to a series of acetates with variable lengths including Z11-18:OAc (cVA), Z11-16:OAc and Z11-14:OAc (Figure 7). On the other hand, OR67d was inactive with a saturated analog (16:OAc), with the isomer (E11-16:OAc), with the analogs with double bond in different positions (Z9-16:OA and Z7-16:OAc), and with the corresponding aldehyde (Z11-16:Ald), alcohol (Z11-16:OH) and acid (Z11-16:COOH) (Figure 7).

**Figure 7.**
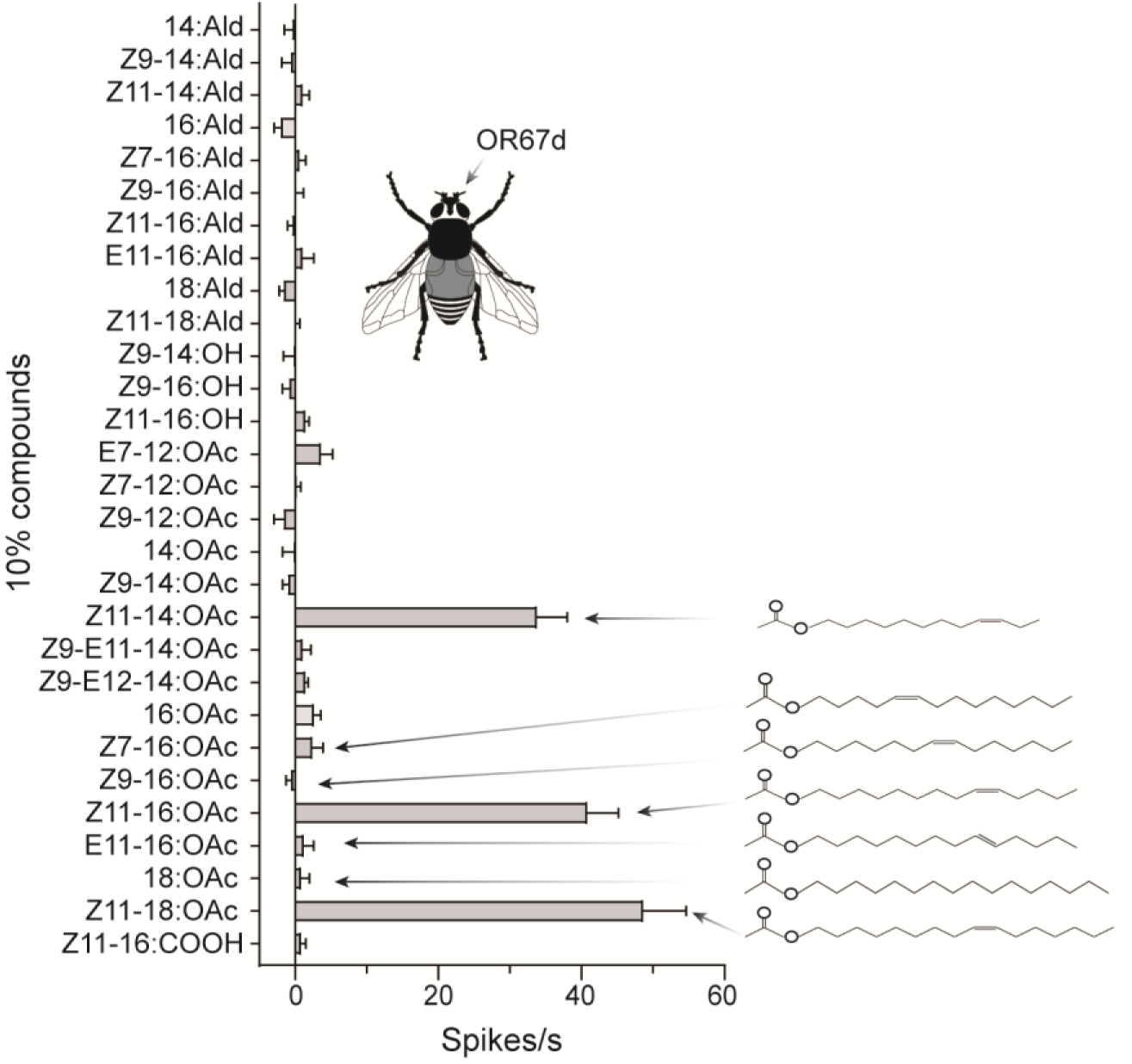
Response spectrum of *Drosophila* OR67d ORN. Recording was made from T1 sensilla of *w*^*1118*^ flies were recoded. OR67d ORN strongly responds to Z11-18:OAc (cVA), Z11-16:OAc, Z11-14:OAc, but not to other structural analogues including E11-16:OAc, Z9-16:OAc, Z11-16:Ald, Z7-16:Ald, Z11-16:OH and Z11-16:COOH. Data are plotted with ± SEM. N = 5 for each recording.

## Discussion

Recently, *Drosophila* mutants lacking OBPs were shown to strongly respond to general odorants, casting a doubt to the necessity of OBPs in olfaction (Xiao et al., 2019). Since PBPs are subtypes of OBPs, the role of PBPs in pheromone detection could also be questioned. In this study, we found that the presence of HarmPBP1 and HarmPBP2 in *Drosophila* T1 sensilla enhanced the sensitivity of HarmOR13 to Z11-16:Ald (Figure 5). This result is in agreement with the defective pheromone sensitivities in several moth PBP mutants (Dong et al., 2019; Shiota et al., 2018; Ye et al., 2017; Zhu et al., 2019). Moreover, *Drosophila* LUSH mutant shows a severe defect in the sensitivity to the major volatile pheromone, cVA (Xu et al., 2005). We may suspect, therefore, although the role of OBPs in general odorants detection could be questioned, PBPs might be important for ensuring a strong sensitivity to pheromone detection.

Another key question is whether PBPs may contribute to the specificity of pheromone detection. In the present work, we observed that HarmPBP1 and HarmPBP2 are colocalized and expressed in the type A sensilla trichodea which house HarmOR13 expressing ORNs. Intriguingly, we also noticed that a small fraction of HarmPBP1 and HarmPBP2 positive cells are not physically associated with HarmOR13 (Fig. 3a, b), hinting at their presences in type B and C sensilla trichodea and their roles in detection of other pheromone components, such as Z9-16:Ald, Z9-14:Ald, Z11-16:OAc. These distribution patterns of HarmPBP1 and HarmPBP2 agree with the their non-selective binding affinities to pheromones (Guo et al., 2012). In addition, the OR67d ORNs of *Drosophila* expressing HarmOR13 are narrowly tuned to the Z11-16:Ald and Z11-14:Ald regardless of the presence of HarmPBP1 or HarmPBP2 (Fig. 6). Recently, mutants of some Lepidoptera lacking PBPs have been reported to show reduced but nondiscriminatory sensitivities to pheromones (Dong et al., 2019; Shiota et al., 2018; Ye et al., 2017; Zhu et al., 2019). For instance, in *S. litura*, knocking-out of the gene encoding *SlitPBP1* had the effect of reducing the antennal response to the three pheromone components, Z9,E11-14:OAc, Z9,E12-14:OAc and Z9-14:OAc by 53%, 60% and 63%. These values were even lower (40%, 43% and 46% respectively) when also the gene encoding *SlitPBP2* was silenced (Zhu et al., 2019). These results indicate reveal only minor effects of PBPs on the pheromone detection. Other studies, instead, suggest more important roles for PBPs. In *Heliothis virescens*, HvirPBP2, but not HvirPBP1 can dramatically sharpen the response breadth of HvirOR13 to Z11-16:Ald [50]. Furthermore, in *B. mori*, BmorPBP1 was shown to strongly imporve the response specificity of BmorOR1 through its binding to bombykol (Große-Wilde et al., 2006). Contradictory findings from studies on different moth species hint at a possibility that species-specific PBPs have different modes of action. However, in the light of the relatively high sequence similarities of orthologue PBPs and the conserved anatomical structure of peripheral olfactory systems across different moth species, the hypothesis of substantial differences in the mode action of PBPs seems unlikely. Nevertheless, our results favor a model in which PBPs do not have a significant effect on the selectivity of ORNs and PRs determine the specificity of pheromones detection. In addition, our work shows the importance of different structural parameters, such as double bonds, functional groups and isomerism for the ligand-receptor affinity (Fig 6, 7). Similar findings also was found in BmorOR1 (Xu et al., 2012). Whether this could be a general phenomenon requires comparative studies on the coding of PRs across different insect species.

In summary, Our study demonstrates that HarmPBP1 and HarmPBP2 contribute to the sensitivity of Z11-16:Ald detection through their similar binding affinities to Z11-16:Ald. Moreover, our study provides a new way to investigate the role of moth PBPs with the advantage of using a live insect expression system.

## Materials and Methods

### Insects

Colonies of *H. armigera* were started from insects collected in the field near Zhengzhou, Henan province in northern China. *H. armigera* were reared under a 16L/8D photoperiod at 26°C and 50% relative humidity. Pupae were sexed and males and females were put into different cages for eclosion. Adults were fed with 10% honey solution daily. For the work relevant to *Drosophila Melanogaster*, an isogenized strain of *w*^*1118*^ was used as a wild-type control for most experiments. *pLUSH-GAL4* fly was obtained from Dean Smith lab at the University of Texas, Southwestern Medical Center (Xu et al., 2005) and *pOrco-LexA::VP16* fly was kindly provided by Tzumin-Lee laboratory on Janelia Research Campus (Lai and Lee, 2006). *Drosophila* lines were maintained under a 16L/8D photoperiod at 26°C and 50% relative humidity and fed by standard yeast molasses food.

### Chemicals

Z11-16:Ald, Z9-16:Ald, Z11-16:OH, Z9-16:OH, Z11-16:OAc, Z9-16:OAc and (Z)-9-tetradecenal (Z9-14:Ald) were purchased from Shin-Etsu Chemical (Tokyo, Japan). Hexadecenal (16:Ald), octadecanal (18:Ald), octadecyl acetate (18:OAc) were purchased from Tokyo Chemical Industry (TCI Shanghai, China). Hexadecyl acetate (16:OAc) was purchased from Toronto Research Chemicals INC (North York, Canada). (Z)-11-hexadecenoic acid (Z11-16:COOH) was purchased from Abcam (Cambridge, UK). Tetradecenal (14:Ald) and tetradecyl acetate (14:OAc) were purchased from Ark Pham (Arlington Heights, IL, USA). (Z)-11-vaccenyl acetate (Z11-18:OAc) was purchased from Cayman Chemical (Ann Arbor, Michigan, USA). (Z)-11-octadecanal (Z11-18:Ald), (E)-11-hexadecenal (E11-16:Ald), (Z)-7-hexadecenal (Z7-16:Ald), (Z)-11-tetradecenal (Z11-14:Ald), (Z)-9-tetradecenol (Z9-14:OH), (E)-11-hexadecenyl acetate (E11-16:OAc), (Z)-7-hexadecenyl acetate (Z7-16:OAc), (Z)-11-tetradecyl acetate (Z11-14:OAc), (Z)-9-tetradecyl acetate (Z9-14:OAc), (Z,E)-9,11-tetradecadienyl acetate (Z9-E11-14:OAc), (Z,E)-9,12-tetradecadienyl acetate (Z9-E12-14:OAc), (Z)-9-dodecenyl acetate (Z9-12:OAc), (Z)-7-dodecenyl acetate (Z7-12:OAc) and (E)-7-dodecenyl acetate (E7-12:OAc) were custom synthesized by Kunbo technology (Kunming, Yunnan, China). The purity was always the highest available (>90%) and was confirmed by gas chromatography-mass spectrometry (GC-MS).

### RNA extraction and first strand cDNA synthesis

Antennae from twenty moths were homogenized in 1 mL QIAzol (Qiagen, Hilden, Germany). RNA was extracted using Qiagen RNeasy Mini kit (Qiagen, Hilden, Germany) and was quantified by Nanodrop 2000 spectrophotometer (Thermo Fisher Scientific Inc., Delaware, USA). The first strand cDNA was synthesized from 2 μg RNA with GoScript™ Reverse Transcriptase (Promega, Madison, WI, USA).

### Preparation of antisera

Rabbit antisera against HarmPBPs and HarmGOBPs were prepared following our published protocol (Guo et al., 2012). The rat antiserum against HarmPBP1 and the mouse antisea against HarmPBP2 and GOBP1 were produced following a standard protocol by Bioss Biotech (Tongzhou, Beijing, China). The mouse antibody against HarmOR13 was custom prepared by Beijing Protein Innovation (BPI, Beijing, China). A peptide of SDGSDLEGVEKVEDI at the intracellular N terminus of HarmOR13 was synthesized and used for antibody production. All the antisera were purified on Protein A column prior to usage.

### Synthesis of probes

A probe of HarmOR13 was synthesized from linearized pGEM-T vector (Promega, Madison, WI, USA). Digoxigenin (DIG)-labeled probes were synthesized with DIG RNA labeling kit version 12 (SP6/T7) (Roche, Mannheim, Germany). Biotin-labeled probes were synthesized by substitution of DIG RNA labeling mix with biotin RNA labeling mix (Roche, Mannheim, Germany). The probes were precipitated by 4 M LiCl and 100% ethanol, followed by a wash with 75% ethanol. Probes for ORs were fragmented into 300 nt snippets by treatment with 80 mM NaHCO_3_, 120 mM Na_2_CO_3_, pH 10.2 at 60 °C. Probes were stored at −80 °C.

### *In situ* hybridization

The procedures for *in situ* hybridization were adapted from Krieger et al. 2002 (Krieger et al., 2004). Briefly, fresh antennae dissected from four-day old male adults were embedded in JUNG tissue freezing medium (Leica Microsystems, Germany). Sections of 12 µm were prepared with a Leica CM 1950 microtome at −22 °C, and mounted on Superfrost Plus Glass Slides (Electron Microscope Science, Hatfield, USA). Slides were dried in air for 10 min, followed by fixing with 4% paraformaldehyde in 0.1 M NaHCO_3_, pH 9.5 at 4 °C for 30 min, then treated with phosphate buffer saline (PBS) (0.85% NaCl, 1.4 mM KH_2_PO4, 8 mM Na_2_HPO_4_, pH 7.1) for 1 min, then 0.2 M HCl for 10 min, and last, washed twice with PBS for 30 sec. After rinsing in 50% deionized formamide (MP Biomedicals, Solon, OH, USA) / 5×SSC (10×SSC: 1.5 M NaCl, 0.15 M Na-citrate, pH 7.0) for 15 min, each slide was treated with 100 µL hybridization buffer (Boster, Wuhan, China) containing DIG-labeled probes. The slides were incubated at 55 °C for 14 h in a humid box wetted with 50% deionized formamide. Subsequently, the slides were washed twice with 0.1×SSC for 30 min at 60 °C, rinsed briefly in TBS (100 mM Tris, pH 7.5, 150 mM NaCl), and incubated in 1% blocking reagent (Roche, Mannheim, Germany) in TBS plus 0.03% Triton-X100 (Merck, Darmstadt, Germany) for 30 min at room temperature. Then, each slide was incubated with 100 µL antidioxigenin alkaline phosphatase-conjugated antibody (Roche, Mannheim, Germany) diluted 1:500 in 1% blocking reagent in TBS plus 0.03% Triton-X100 at 37 °C for 1 h. After three washes for 5 min with TBS plus 0.05% Tween-20 and a short rinse in DAP buffer (100 mM Tris, pH 9.5, 100 mM NaCl, 50 mM MgCl_2_), DIG-labeled probes were visualized by HNPP/Fast Red (Roche) and Biotin-labeled probes were visualized by TSA kit (Perkin Elmer, MA, USA) including an anti-biotin streptavidin horse radish peroxidase-conjugate and fluorescein-tyramides as a substrate. Pictures were taken with a Zeiss LSM 710 confocal microscope (Zeiss, Oberkochen, Germany).

### Immunostaining

Fresh antennae were fixed in 4% paraformaldehyde for 2 hours and treated with 25% sucrose in 0.1 M NaHCO3, pH 7.4 at 4°C overnight. The antennae were embedded in JUNG tissue freezing medium (Leica Microsystems, Germany) and sliced into 12 µm sections on a Leica CM 1950 microtome. The sections were mounted onto adhesion microscope slides (CITOGLAS, Haimen, Jiangsu, China) and dried for 1 h at room temperature. The sections were treated with 0.5% Triton X-100 (Sigma-Aldrich, USA) and gently washed three times with 0.1% Saponin (Sigma-Aldrich, USA) in PBS, pH 7.4, followed by blocking with goat serum (Solarbio Biotech, Beijing, China) for 30 min at room temperature. Then, the samples were incubated with diluted antisera (1:500 dilution for anti-PBPs and anti-GOBPs sera; 1:150 for anti-HarmOR13 serum) at room temperature for 2 h and subsequently at 4°C overnight. The slides were washed four times for 5 min each and treated with 1:1000 diluted Alexa Fluor® conjugated sec antibody (Cell Signaling Technology, Danvers, MA, USA) for 1 h at room temperature. Finally, after 4 times of washing, the slides were mounted in anti-fade mountant (Beyotime Tech, Beijing, China). Confocal images were taken using Zeiss LSM 710 confocal microscope (Zeiss, Oberkochen, Germany).

### Transgenic flies

The full-length coding DNA sequences (CDS) of *HarmPBP1, HarmPBP2* and *HarmPBP3* were cloned into p10 plasmid (pJFRC-28-10-10×UAS-IVS-GFP-P10, Addgene plasmid # 36431). For phiC31 integrase-mediated P-element insertion on chromosome 2, p10 plasmids were injected into attp40 fly embryos (*y[1] v[1] P{y[*+*t7*.*7]=nos-phiC31\int*.*NLS}X;P{y[*+*t7*.*7]=CaryP}attP40*, BDSC # 25709) by Fungene Biotechnology (Qidong, Jiangsu, China) to generate *UAS-HarmPBPs* transgenic flies for further crossings. Female or male *CyO/Sp;LUSH-GAL4* flies were crossed with female or male *UAS-HarmPBP;Sb/TM6B* to generate *UAS-HarmPBP/CyO*; *LUSH-GAL4/TM6B* flies. Finally, *UAS-HarmPBP;LUSH-GAL4* homozygotes were obtained by sibling crosses and maintained as stable lines for further experiments. For generation of *LexAop-HarmOR13* transgenic flies, the full length of CDS was cloned into p10 plasmid in which the *UAS* sequence was replaced with *LexAop* sequence. Following the same injection and cross procedures, the *LexAop-HarmOR13;ORco-LexA* was established as a stable stock for following experiments.

### Single sensillum recording

Single sensillum recordings were performed as previously described (Guo et al., 2020). For SSRs, 30 μl of diluted odorant in paraffin oil was placed on a small piece of filter paper (1.5 cm^2^) which was inserted into a 5.75-inch Pasteur pipette (ANPEL Lab Tech, Shanghai, China). A constant stream (30 ml/s) of air passed over an activated charcoal filter and then was humidified prior to passing over the preparation. The stimulation pulse was 300 ms in all recordings. To avoid sensory adaptation during recordings, the stimuli were presented at random with an interval of at least 1 min.

### Analysis of data

The two-tailed Student’s *t* test and two-way ANOVA with *post hoc* Tukey’s multiple comparisons was performed with Graphpad Prism 8 software (GraphPad Software, San Diego, CA, USA).

## Acknowledgements

We are grateful to Tzumin-Lee laboratory at Janelia Research Campus for kindly providing the *pOrco-LexA* fly and to Dean Smith laboratory at the University of Texas, Southwestern Medical Center for sharing *pLUSH-GAL4* stock. We owe great thanks to Dr. Paolo Pelosi of the Austrian Institute of Technology GmbH, Biosensor Technologies for his critical comments on a draft of this manuscript and to Dr. Jürgen Krieger for his invaluable suggestions to set up *in situ* hybridization experiments. We are greatly indebted to Chuan Zhou laboratory in the Institute of Zoology, Chinese Academy of Sciences for providing fly food. This work was funded by the National Natural Science Foundation of China (Grant No.31801748 and 31772528).

---

**Figure S1.**
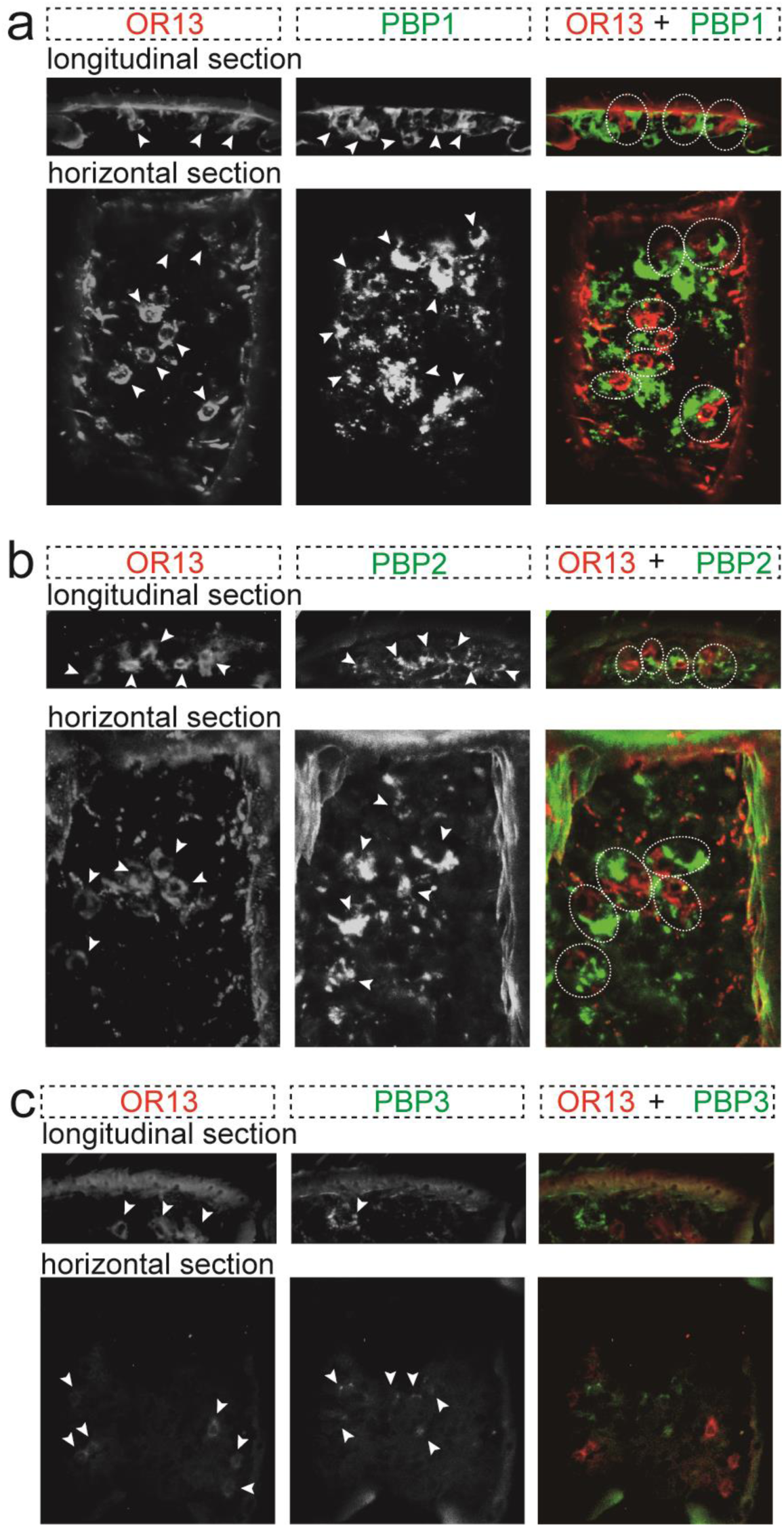
Two-color fluorescence *in situ* hybridization to localize *HarmOR13* mRNA and *HarmPBPs* mRNA. Close localizations are highlighted by dash circles. **(a)** *HarmOR13* expressing ORNs are surrounded by *HarmPBP1* expressing supporting cells. **(b)** *HarmOR13* and *HarmPBP2* are adjacently expressed. **(c)** *HarmOR13* mRNA is physically separated from *HarmPBP3* mRNA. Scale bars: 20 µm.

